# Dissecting the roles of local packing density and longer-range effects in protein sequence evolution

**DOI:** 10.1101/023499

**Authors:** Amir Shahmoradi, Claus O. Wilke

## Abstract

What are the structural determinants of protein sequence evolution? A number of site-specific structural characteristics have been proposed, most of which are broadly related to either the density of contacts or the solvent accessibility of individual residues. Most importantly, there has been disagreement in the literature over the relative importance of solvent accessibility and local packing density for explaining site-specific sequence variability in proteins. We show here that this discussion has been confounded by the definition of local packing density. The most commonly used measures of local packing, such as the contact number and the weighted contact number, represent by definition the combined effects of local packing density and longer-range effects. As an alternative, we here propose a truly local measure of packing density around a single residue, based on the Voronoi cell volume. We show that the Voronoi cell volume, when calculated relative to the geometric center of amino-acid side chains, behaves nearly identically to the relative solvent accessibility, and both can explain, on average, approximately 34% of the site-specific variation in evolutionary rate in a data set of 209 enzymes. An additional 10% of variation can be explained by non-local effects that are captured in the weighted contact number. Consequently, evolutionary variation at a site is determined by the combined action of the immediate amino-acid neighbors of that site and of effects mediated by more distant amino acids. We conclude that instead of contrasting solvent accessibility and local packing density, future research should emphasize the relative importance of immediate contacts and longer-range effects on evolutionary variation.

## Introduction

A variety of site-specific structural characteristics have been proposed over the past decade to predict protein sequence evolution from structural properties. Among the most important and widely discussed are the Relative Solvent Accessibility (RSA)^1–15^, Contact Number (CN)^11–13,16–21^, measures of thermodynamic stability changes due to mutations at individual sites in proteins^3,22–25^, protein designability^26,27^, and measures of local flexibility, such as the B factor^13,18,28^, or flexibility measures based on elastic network models^20,29^ and Molecular Dynamics (MD) simulations^13^.

It is conceivable that the majority of these site-specific quantities predicting evolutionary rate (ER) simply act as different proxy measures of an underlying biophysical constraint, such as the strength of amino acid interactions at individual sites. Indeed, Huang et al. ^20^ recently proposed that the underlying constraint is mechanistic stress, which can be estimated by calculating the weighted contact number (WCN) at each site. Quantities such as relative solvent accessibility and local flexibility are expected to correlate with WCN, because all these measures depend to some extent on the density of contacts around an amino acid. Huang and coworkers thus argued that the *local packing density* is the primary constraint of protein sequence evolution^11,12,20^.

However, a priori it is not clear whether quantities based on the contact-number concept (either CN or WCN) are truly measures of *local* packing density. All these measures involve adjustable free parameters in their definitions, and depending on the parameter choices, the measures can incorporate substantial long-range effects. In particular, WCN by default uses an inverse-square weighting, which assumes that amino acids at an arbitrary distance away from the focal residue still exert some measurable effect on that residue. Thus, some of the best predictors of evolutionary variation seem to confound local and longer-range effects, and the relative importance of the immediate neighborhood of an amino acid vs. longer range effects is not known.

To separate out the effects of direct-neighbor and longer-range effects, we here derive a new set of structural characteristics that, unlike CN and WCN, depend only on the immediate contacts of a given amino acid and do not involve freely adjustable parameters in their definitions. We achieve this goal by employing tessellation methods developed in the field of computational geometry. We find that quantities based on tessellation methods explain much but not all of the variance in ER that WCN captures. Therefore, we conclude that longer-range effects beyond the first coordination shell play a significant role in protein sequence evolution.

## Methods

### Protein data set and site-specific sequence variability

We analyzed the same data set of 209 monomeric enzymes we have previously analyzed^24^ and that was originally published with four additional proteins^11^. The enzymes were originally randomly picked from the Catalytic Site Atlas 2.2.11^30^, range from 95 to 1287 amino acids in length, and include representatives from all six main EC functional classes ^31^ and domains of all main SCOP structural classes^32^. Evolutionary rates at all sites in all proteins were calculated as described^24^. In brief, we employed the software Rate4Site^33^, using the empirical Bayesian method and the amino-acid Jukes-Cantor mutational model.

### Packing density and relative solvent accessibility

Local Packing Density (LPD) for individual sites in all proteins was calculated according to the two most-commonly used definitions, Contact Number (CN) and Weighted Contact Number (WCN) (defined in Eqs. 2 and 4 in Results). Both CN and WCN generally require each residue to be represented by a single point in space (but see the work by Marcos and Echave^21^, who used a two-point representation). Here, we examined seven different ways to choose the reference point for each amino acid: We considered all backbone atomic coordinates, i.e., the coordinates of N, C, O, or C_*α*_, the coordinates of the first heavy atom in the side-chains, C_*β*_, and two more representations obtained by averaging over the coordinates of all heavy atoms in the amino acid including the backbone atoms (denoted as AA coordinates) or of all side-chain heavy atoms (denoted as SC coordinates). For each of these seven representations, we calculated CN and WCN for each residue in each protein. Appendix A shows that the SC representation results in the highest correlations between structural measures and sequence evolution, and therefore we used SC coordinates for all results reported in the main body of the manuscript.

We used the DSSP software^34^ to calculate the Accessible Surface Area (ASA) for each residue in each protein, and we normalized these ASA values by their theoretical maximum^35^ to obtain Relative Solvent Accessibility (RSA).

### Voronoi tessellation

We used the VORO++ software^36^ to calculate the relevant Voronoi cell properties of all sites in all proteins in the dataset. Among the most important properties are the length of the cell edges, surface area and volume, number of faces of each cell, and the cell eccentricity, defined as the distance between the cell’s seed and the geometrical center of the cell. In addition, the cell *sphericity*, which is a measure of the compactness of the cell, can be defined as

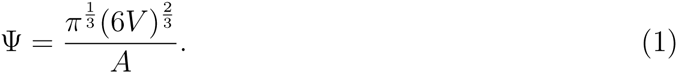

Here, *V* and *A* represent the volume and the area of the cell, respectively. By definition, Ψ falls between 0 and 1. The limit of Ψ = 1 indicates that the cell is perfectly spherical, while the limit of Ψ = 0 indicates that the cell is a 2-dimensional object that has no volume but only surface area.

Each Voronoi cell is a polyhedron consisting of a set of polygons joined together at their edges. The total edge length of a cell was calculated as the sum of the edge lengths of all such polygons. The resulting sum was divided by two to obtain the total edge length, since each edge is shared between two polygons. Similarly, the cell surface area was calculated as the sum of the surface areas of all polygons that bounded it. Finally, the cell volume was defined as the volume of the Voronoi polyhedron.

As for CN and WCN, we considered seven different coordinate representations, and we found that the SC coordinates performed best (Appendix A). Therefore, unless otherwise noted, all results reported in the main body of the text are based on SC coordinates.

### Edge effects in Voronoi partitioning of protein structures

One potential caveat with Voronoi tessellation of finite structures is the existence of *edge effects*. Sites that are close to the surface of the protein can be associated with Voronoi cells that are open, only bounded by the cubic box containing the protein. We performed the following analysis to verify that these edge effects did not influence the observed sequence-structure correlations. We first identified all open cells in all proteins by examining the fraction of change in the cell volumes upon changing the size of the cubic box that contained the protein. For each protein, once the open cells were identified, we multiplied their volumes by a normalizing factor such that the smallest-volume open cell had a larger volume than largest closed cell. We then ordered the open cells by the fraction of volume change observed upon changing the box size. In other words, the more the cell volume increased upon increasing the box size, the larger the volume of that cell was assumed to be, compared to all other open cells in the protein. Finally, we calculated Spearman’s correlation coefficient between this ordered set of cell volumes (that included both open and closed cells) and the sequence evolutionary rates. We then compared the computed correlation coefficients to those obtained by using the original cell volumes without corrections for edge effects. The resulting distributions of correlation coefficients for the original and corrected cell volumes were nearly identical (mean difference < 0.001).

For our dataset of 209 monomeric proteins, we concluded that edge effects due to Voronoi tessellation appear to have virtually no influence on the observed sequence-structure correlations. We reached similar conclusions when open cells were alternatively ranked by other criteria, such as the fractional changes in cell area upon changing the box size. Thus, the Voronoi cell characteristics, in particular cell volume and cell area, can be safely used in predicting sequence variability, as long as a sufficiently large enclosing box is used to calculate the Voronoi cell characteristics of open cells.

One exception, however, is cell sphericity, as defined in Eqn. 1. This quantity behaves differently for open and closed cells. Unlike the case for closed cells in the protein’s interior, the sphericity of open cells on the surface of the protein is generally *positively* correlated with site-specific evolutionary rates. This is an artifact of the edge effects in the tessellation of finite structure of protein. A possible correction to the sphericity of open cells could be therefore applied by negating the value of sphericity for open cells. This correction resulted on average in a 0.05 increase in the correlation coefficients of sphericity with ER. However, since this correction has no obvious justification based on first principles, we did not employ it for further analyses. More importantly, the inclusion or exclusion of edge-effect corrections to cell sphericity did not substantively affect any of the major findings of this work.

### Statistical analysis and data availability

Statistical analyses were carried out with the software package R^37^. We used Spearman correlations and partial Spearman correlations throughout this work. Partial Spearman correlations were calculated using the R package *ppcor*^38^.

All data and analysis scripts necessary to reproduce this work are publicly available to view and download at https://github.com/shahmoradi/cordiv.

## Results

### Optimal definition of contact number incorporates longer-range effects

Prior work on the same enzyme data set that we have analyzed here has shown that contact number generally outperforms all other structural predictors of site variability^11,12,20^. This finding has been interpreted as indicating that local packing density is the primary evolutionary constraint imposed by protein structure. However, to what extent the contact number (CN) is actually a truly local measure is unclear. The contact number at a site *i*, CN_*i*_, is defined as the number of amino acids within a fixed radius *r*_0_ from that site,

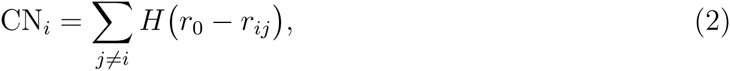

where *r*_*ij*_ is the distance between sites *i* and *j* and *H*(*r*) is the Heaviside step function

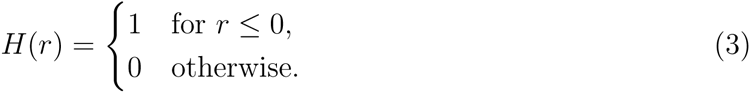

The location of individual sites is commonly chosen as the position of the residues’ C_*α*_ backbone atoms, but recent work has shown that the geometric center of the side chain may be a better choice^21^. Irrespective of the particular choice of reference point for each site, how local of a measure CN_*i*_ actually is depends on the arbitrary cutoff parameter *r*_0_. There is no consensus on the optimal value of this cutoff distance, although it is typically chosen in the range of 5Å to 18Å^6,12,39,40^. While a cutoff of 5Å likely captures only immediate neighbors of the focal amino acid, a cutoff of 18Å will capture many amino acids that do not directly interact with the focal amino acid, and thus captures both local and non-local packing-density information.

To determine at what cutoff parameter *r*_0_ the contact number had the most predictive power for evolutionary variation, we calculated, for each protein in our data set, the Spearman correlation between evolutionary rate (ER) and contact number CN across the protein as a function of *r*_0_ (Figure 1A). (A similar analysis has previously been published^12^.) We found that, on average, the correlation started to rise rapidly around 5Å, reached a maximum around 14.3Å, and then started to decline slowly for larger *r*_0_ (see also Table 1). Since the maximum correlation was located at a cutoff value that captured at least two complete shells of amino acids around the focal residue, this analysis suggested that non-local effects play some role in constraining amino-acid variability.

**Table 1:** The first, second (median), and the third quartiles of the distribution of the best free parameters of the Contact Number (CN) and the Weighted Contact Number (WCN) resulting in the strongest correlations with sequence evolutionary rates. The best parameter distributions are obtained by tuning the free parameters of CN and WCN for individual proteins in the dataset of 209 monomeric enzymes such that the strongest correlation with evolutionary rates is obtained.

| Local Packing Density Measure | Best free-parameter |  |  |
| --- | --- | --- | --- |
|  | 25% quartile | median | 75% quartile |
| CN | $r_0 = 9.3 \text{ [\AA]}$ | $r_0 = 14.3 \text{ [\AA]}$ | $r_0 = 19.6 \text{ [\AA]}$ |
| WCN | $\alpha = -2.7$ | $\alpha = -2.3$ | $\alpha = -1.5$ |

**Figure 1:**
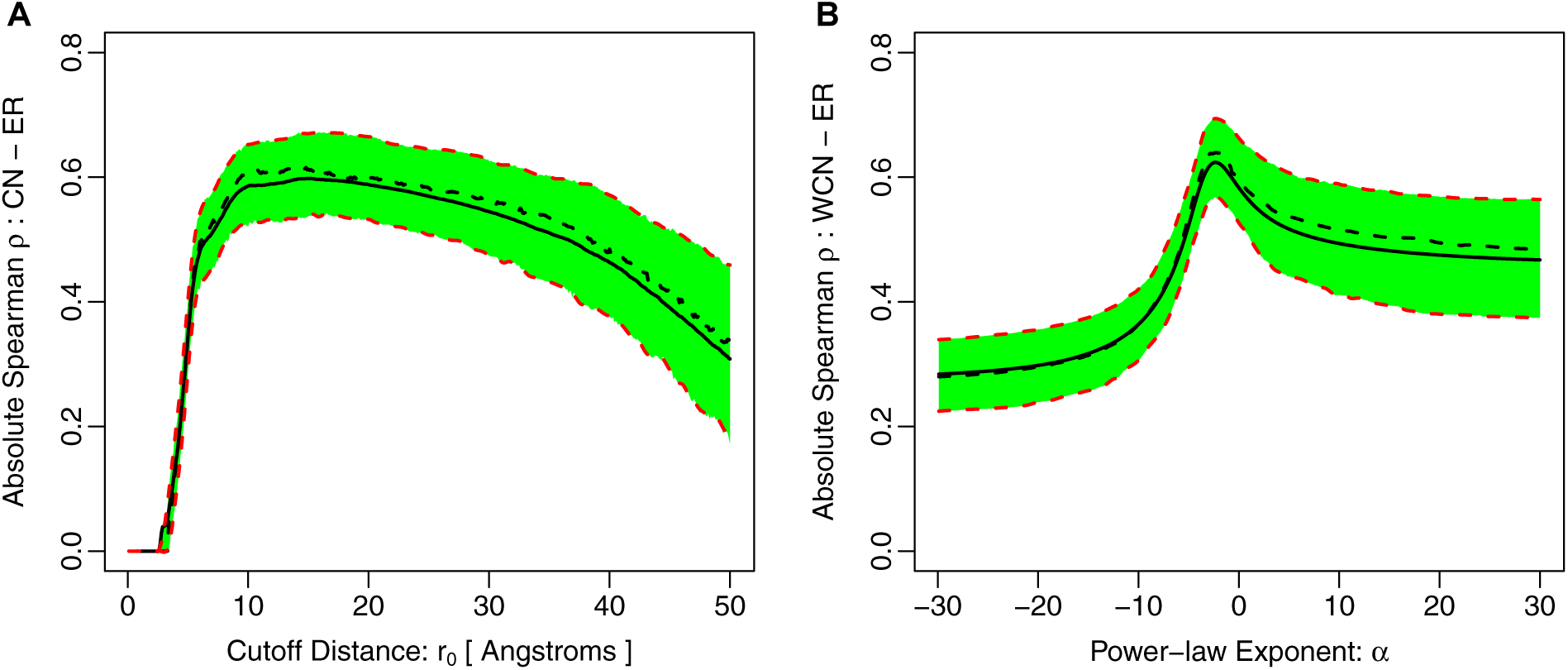
Absolute correlation of evolutionary rate with (A) Contact Number and (B) Weighted Contact Number for varying degrees of locality of these quantities. In (A), we vary the cutoff parameter *r*_0_ in Eq. 2 from 0 to 50Å. In (B), we vary the exponent *α* in Eq. 4 from −30 to 30. In each plot, the solid black line represents the mean correlation strength in the entire dataset of 209 proteins and the dashed black line indicates the median of the distribution. The green-shaded region together with the red-dashed lines represent the 25% and 75% quartiles of the correlation-strength distribution. Note that for the case of WCN with *α* < 0 the sign of the correlation strength *ρ* is the opposite of the sign of *ρ* with *α* < 0. In addition, *ρ* is undefined at *α* = 0 and not shown in this plot. The parameter values at which the correlation coefficient reaches the maximum over the entire dataset are given in Table 1.

The arbitrariness of the cutoff *r*_0_ in the CN definition has been long recognized, and therefore some authors^39^ have suggested an alternative quantity known as the Weighted Contact Number (WCN). For a given site *i* in a protein of length *N*, WCN_*i*_ is defined as the sum of the inverse-squared distances to all other sites in protein,

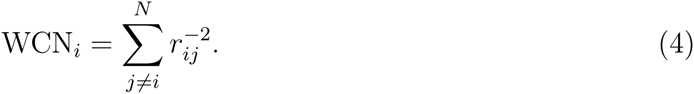

It turns out that WCN generally performs better than CN at predicting ER^11,12,20^. However, WCN is clearly *not* a local measure. By virtue of the exponent −2, the aggregated influence of all amino acids at distance *r* on WCN is the same, irrespective of *r*. In other words, the factor 1*/r*^2^ exactly compensates the geometrical increase in the number of residues in each shell, for arbitrarily large radii *r*. Thus, rather than just measuring local packing density, WCN also measures where the focal amino acid falls relative to the overall shape and distribution of amino acids in the protein. For example, in a perfectly spherical protein of uniform density, WCN would be a direct proxy of the distance to the geometric center of the protein.

Moreover, just as was the case with CN, the definition of WCN contains a somewhat arbitrary parameter^41^, the exponent −2. If we chose a more negative exponent, then WCN would be a more local measure of packing density, putting less weight on increasingly distant amino acids. Likewise, if we chose a less negative exponent, WCN would put more weight on more distant amino acids and less weight on amino acids close in. Therefore, just as we had done with CN, we calculated the correlations between WCN and ER as a function of the exponent *α* (Figure 1B). We found that the correlations peaked at *α ∼* −2.3, almost exactly the canonical value of −2.

Evidently, both CN and WCN perform best at describing ER variation when they capture a non-negligible contribution of longer-range effects, beyond the immediate neighborhood of each focal site. To disentangle the contributions of local and longer-range effects to sequence evolution, we next proceeded to develop a measure of packing density that by definition depended on only the immediate neighbors around a single amino acid.

### Measures of local packing density based on Voronoi tessellations

To describe the local surroundings of a residue inside a protein in an unambiguous manner, we need to partition the space inside a protein into regions that uniquely belong to specific residues. Such structural partitioning has a long history in the analysis of protein structures^42,43^. In particular, the Voronoi tessellation and its dual graph, the Delaunay triangulation, have been used in several studies analyzing the internal structure of proteins and/or developing empirical potentials ^44–46^. The Voronoi tessellation divides the Euclidean space into regions, called *cells*, and each belonging to exactly one centroid point (seed), such that the cell corresponding to each centroid point consists of every region in space whose distance is less than or equal to its distance to any other centroid point (Figure 2).

**Figure 2:**
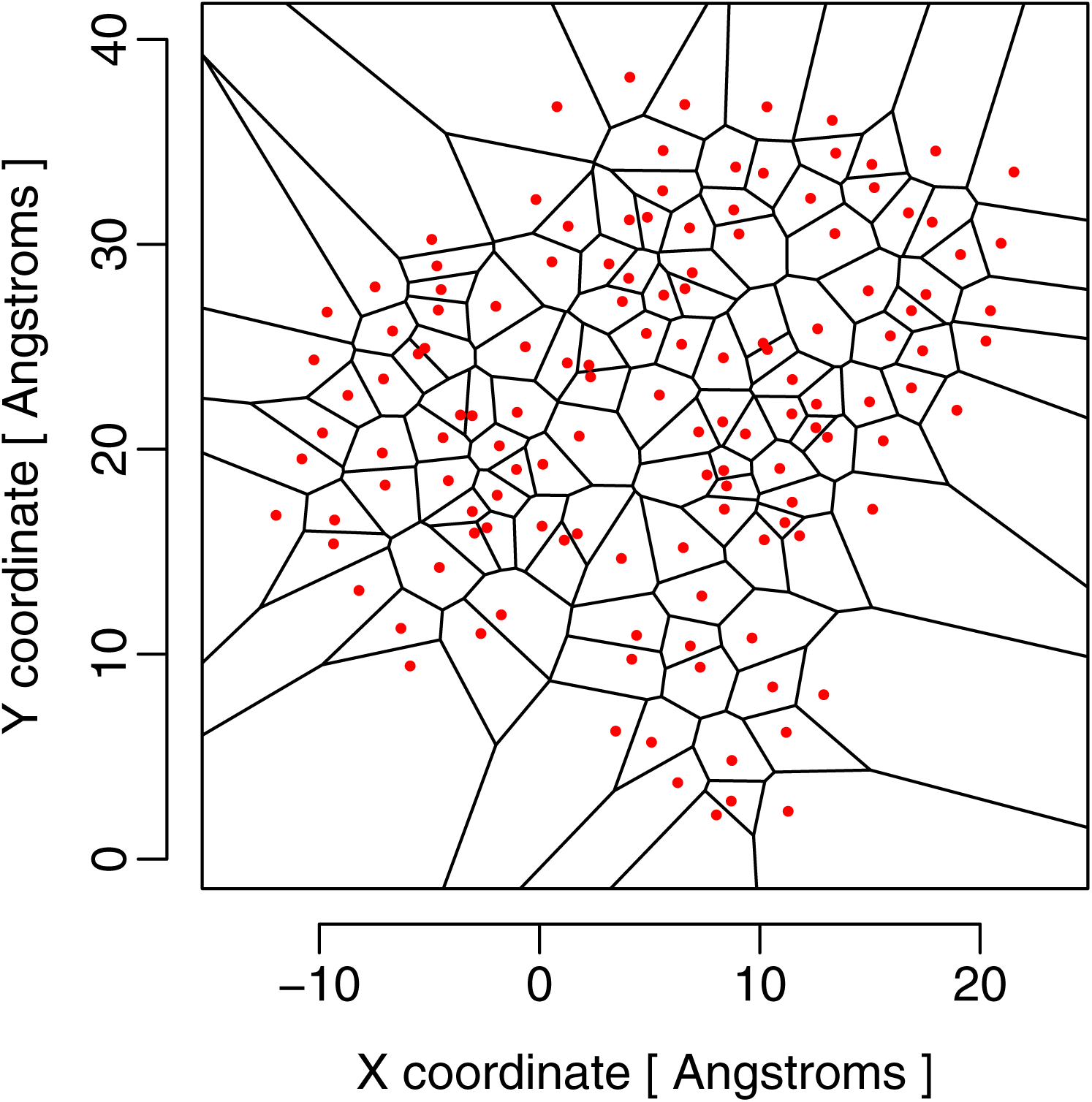
Example of a Voronoi tessellation in two dimensions. The red dots represent the seed points, and the black lines delineate the Voronoi cells. For protein structures, the tessellation is carried out in three dimensions.

When applying a Voronoi tessellation to a protein structure, we usually choose one seed point per residue in the structure. As is the case with CN and WCN, there are multiple possible choices for which point best represents each residue. While many authors simply use the locations of the C_*α*_ backbone atoms, we can also use the geometric center of the amino-acid side-chain, the geometric center of the entire amino acid, or any particular heavy atom in the amino-acid backbone or side-chain. We show in Appendix A that we obtain the strongest correlations of Voronoi cell properties with site-specific sequence variability of proteins if we use the geometric centers of the residue side chains as Voronoi seeds. Thus, we used these specific seeds for the remainder of this work.

How can the Voronoi tessellation be used to describe the local packing density in a protein? Most importantly, the volume of a Voronoi cell is a direct measure of the amount of space surrounding a particular residue. More densely packed regions of the protein will yield smaller cell volumes than less densely packed regions. However, in addition to cell volume, several other cell properties also measure the local geometric arrangement of residues surrounding the focal one, such as cell edge length, cell area, cell eccentricity, and cell sphericity (see Methods for definitions of these quantities). We asked which of these quantities could be used to predict ER, and we found that they all did (Figure 3A). Specifically, we found that cell volume and surface area were most strongly correlated with ER, followed by the cell eccentricity, total edge length, and the cell’s sphericity (Figure 3A and Table 3). All of the Voronoi cell characteristics except sphericity correlated positively with ER, while sphericity displayed negative correlations.

**Table 2:** The percentage of variance of site-specific evolutionary rates explained by Voronoi cell volume (representing the inverse of site-specific packing density) and Weighted Contact Number controlling for cell volume (representing longer-range effects). In each case, the first, second (median) and the third quartiles of the percentage distribution are reported.

| Structural Quantity | Percentage of ER Variance Explained |  |  |
| --- | --- | --- | --- |
|  | 25% quartile | median | 75% quartile |
| Voronoi Cell Volume | 27% | 34% | 41% |
| WCN, controlling for Cell Volume | 5% | 10% | 16% |

**Table 3:** *P*-values of pairwise paired t-tests of the absolute correlation strengths of evolutionary rates (ER) with different Voronoi cell characteristics: volume, area, total edge length, sphericity, eccentricity. Cell area and cell volume perform equally well in predicting ER, while all other cell characteristics perform significantly worse (Fig. 3). The *P*-values were corrected for multiple testing using the Hochberg correction^52^.

|  | ER – Area | ER – Eccentricity | ER – Edge | ER – Sphericity |
| --- | --- | --- | --- | --- |
| ER – Volume | $6 \times 10^{-20}$ | $10^{-26}$ | $2 \times 10^{-21}$ | $1 \times 10^{-60}$ |
| ER – Sphericity | $2 \times 10^{-70}$ | $2 \times 10^{-43}$ | $4 \times 10^{-88}$ | – |
| ER – Edge | $5 \times 10^{-86}$ | $8 \times 10^{-15}$ | – | – |
| ER – Eccentricity | $6 \times 10^{-37}$ | – | – | – |

**Figure 3:**
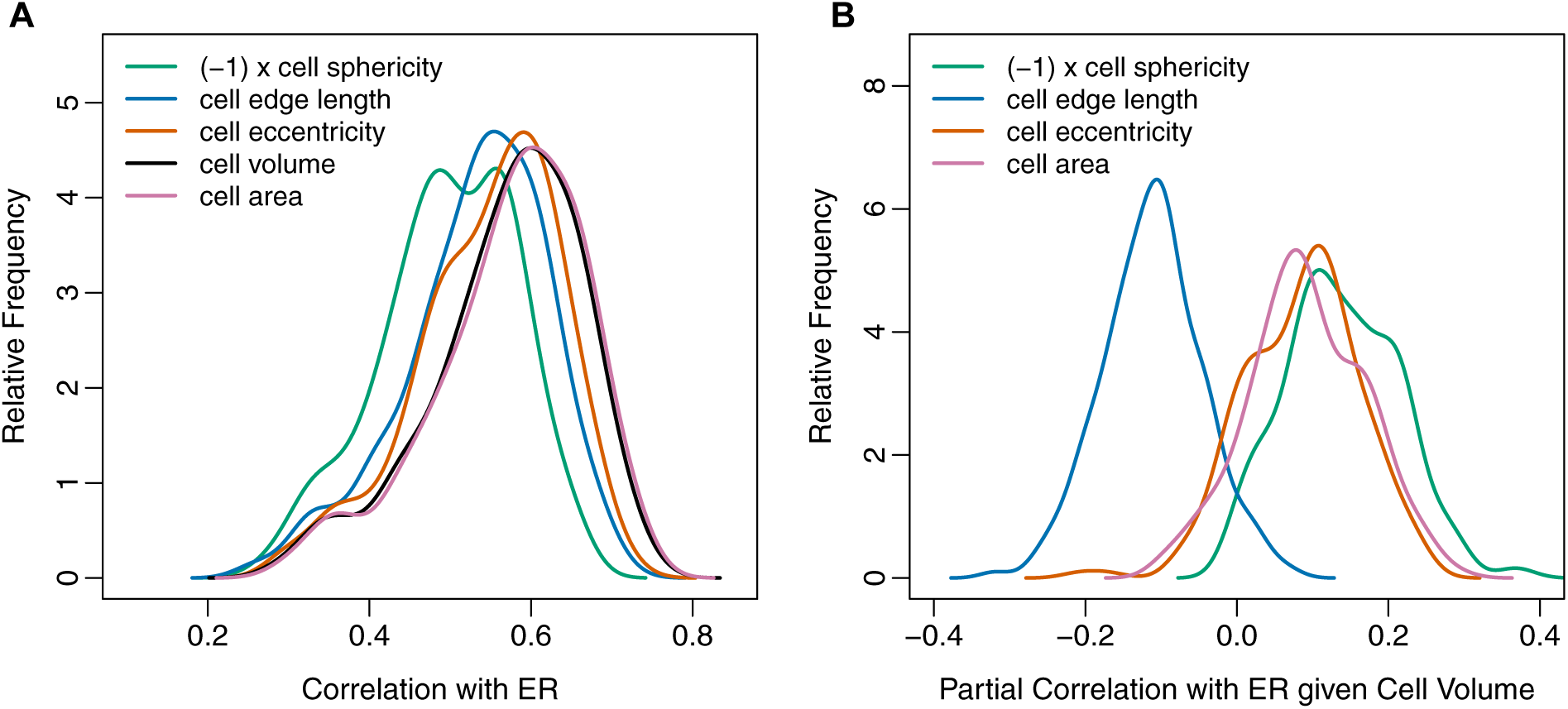
Correlations and partial correlations of ER with various Voronoi cell properties. (A) Distributions of ER correlations with Voronoi cell properties sphericity, edge length, eccentricity, volume, and area. Note that all cell characteristics correlate positively with ER, except sphericity which correlates negatively with ER. We show here the correlations of ER with (*-*1) *×*cell sphericity so that the correlation distributions all appear on the same positive scale. The correlations for cell area and cell volume are not significantly different from each other, and are higher than the correlations for all other Voronoi cell properties (Table 3). (B) Distributions of partial correlations of ER with Voronoi cell properties, controlling for cell volume. All partial correlations are relatively small, with medians of approximately 0.1. Therefore, none of these cell properties provide much independent information about ER once cell volume is accounted for.

In this context, we would like to emphasize that most Voronoi cell characteristics increase with decreasing local packing density. For example, the larger the cell volume or cell surface area, the fewer amino acids are located in the immediate neighborhood of the focal residue. By contrast, increasing sphericity implies increasing packing density. In general, a polyhedron with a higher number of faces corresponds to higher sphericity, the limiting case of which is a perfect sphere with an infinite number of faces. In the context of Voronoi polyhedra in proteins, a higher number of faces of a cell indicates more amino-acid neighbors around the site of interest, and therefore more interactions among them. Consequently, correlations with cell sphericity tend to have the opposite sign from correlations with other Voronoi cell characteristics.

Not unexpectedly, we found that the Voronoi cell characteristics correlated strongly with each other. Therefore, we next used cell volume (measuring inverse local packing density) as the reference and calculated the partial correlations of ER with Voronoi cell properties controlling for cell volume. We found that these partial correlations were generally quite low, with median absolute values of approximately 0.1 (Figure 3B). Cell edge length and cell sphericity displayed negative partial correlations with ER while cell eccentricity and cell area displayed positive partial correlations. In conclusion, with regards to ER variation, the cell area, volume, and edge length appear to represent largely the same underlying property of a Voronoi cell, its size.

### Influence of longer-range effects on sequence evolution

We next asked how Voronoi cell volume compared as predictor of ER relative to the more commonly studied quantities WCN and RSA. As seen in Figure 4A, Voronoi cell volume, RSA, and C_*α*_ WCN all displayed comparable correlations with ER, with median (absolute) correlation coefficients of 0.59, 0.57, and 0.56, respectively. By contrast, side-chain WCN out-performed these predictors by a substantial margin, with median (absolute) correlation coefficient of 0.64 (see Table 4 for pairwise paired t-tests among all distributions). The superior performance of side-chain WCN on this data set is consistent with previous independent results in the literature^21^.

**Table 4:** *P*-values of pairwise paired t-tests of the absolute correlation strengths of evolutionary rates (ER) with four structural characteristics depicted in Figure 4A: WCN (SC), WCN (CA), cell volume, and RSA. The *P*-values were corrected for multiple testing using the Hochberg correction^52^.

|  | ER – WCN(SC) | ER – WCN(CA) | ER – Volume |
| --- | --- | --- | --- |
| ER – RSA | $2 \times 10^{-12}$ | 0.083 | $6 \times 10^{-17}$ |
| ER – Volume | $7 \times 10^{-6}$ | $5 \times 10^{-4}$ | – |
| ER – WCN(CA) | $5 \times 10^{-11}$ | – | – |

**Figure 4:**
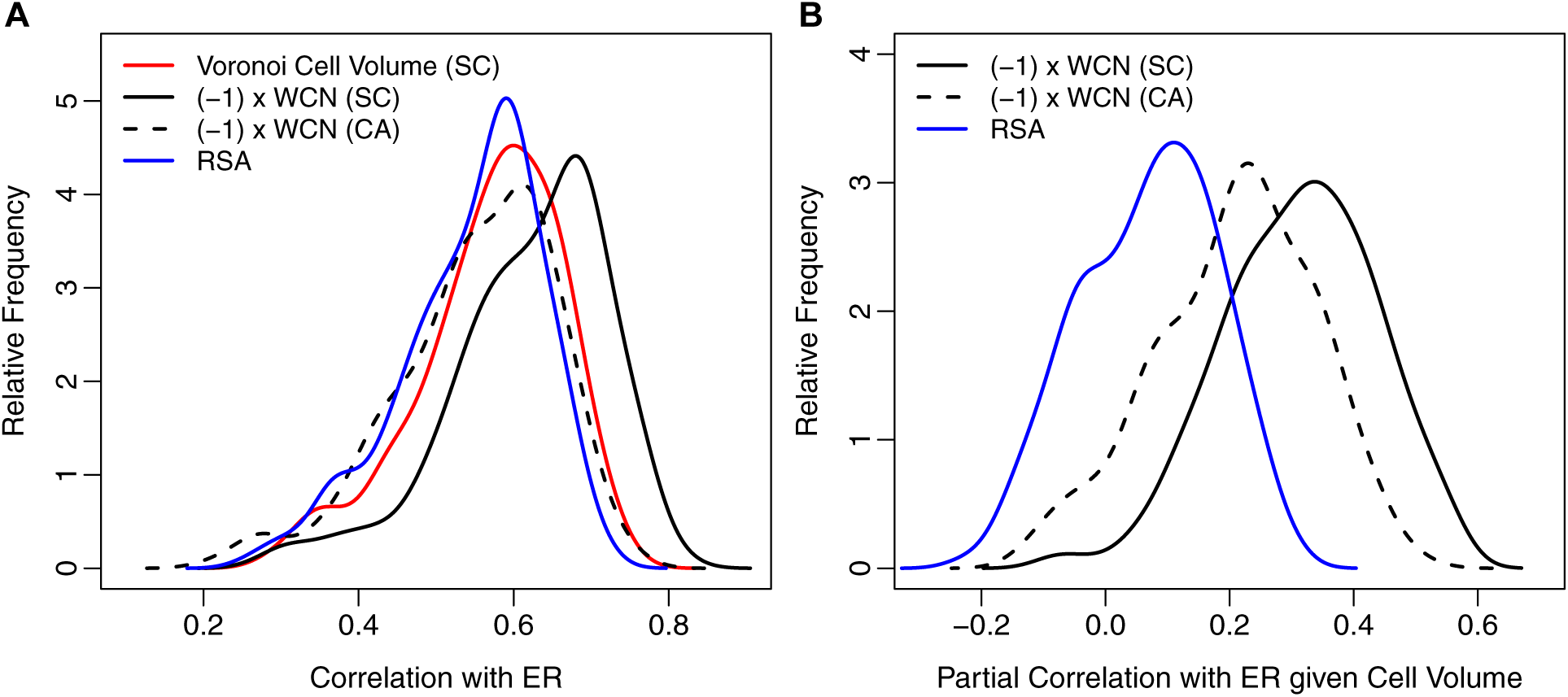
Correlations and partial correlations of ER with Voronoi cell volume, WCN, and RSA. (A) Distributions of ER correlations with Voronoi cell volume, WCN (using side-chain and C_*α*_ coordinates, denoted by SC and CA, respectively), and RSA. Note that the WCN measures correlate negatively with ER. We show here correlations of ER with (-1) *×*WCN so that the correlation distributions all appear on the same positive scale. The correlations of ER with WCN (SC) are significantly higher than the correlations of ER with all other quantities shown (Table 4). (B) Distributions of partial correlations of ER with WCN and RSA, controlling for cell volume. The partial correlation of ER with −WCN are substantial, with median values of 0.32 (side-chain WCN) and 0.21 (C_*α*_ WCN). By contrast, the partial correlations of ER with RSA largely vanish, with a median value of 0.09.

Note that both Voronoi cell volume and RSA are purely local quantities, determined entirely by the immediate neighborhood of an amino acid. In fact, they correlate strongly with each other, and measure approximately the same quantity (Appendix A). By contrast, as mentioned previously, WCN takes into account the amino-acid arrangement in the entire protein, and it equally weighs each shell with radius *r* around the focal residue. Thus, to disentangle local from non-local effects, we calculated partial correlations of ER with WCN while controlling for Voronoi cell volume. In this way, we could quantify the residual influence of WCN—that is, the influence of longer-range effects beyond immediate neighbors—on sequence variability. As a control, we performed the same calculation for RSA instead of WCN.

We found that side-chain WCN displayed substantial residual correlations with ER after controlling for Voronoi cell volume (Figure 4B). The distribution of (absolute) partial correlation coefficients had a median value of 0.32, with an interquartile range of 0.17 (from 0.23 to 0.40). Thus, after controlling for local packing, the longer-range effects appeared to explain approximately 10% of the site-specific sequence evolutionary rates. Similarly, there were significant residual correlations of C_*α*_ WCN with ER after controlling for Voronoi cell volume, with a median (absolute) correlation coefficient of 0.21. By contrast, the residual correlations of RSA with ER were by-and-large negligible. The median was only 0.09, with an interquartile range of 0.17 (from −0.02 to 0.15).

Thus, in conclusion, either WCN definition (side-chain or C_*α*_) provided a significant, independent contribution to explaining ER when purely local effects of packing were controlled for by Voronoi cell volume. By contrast, RSA did not make an independent contribution once Voronoi cell volume was controlled for, as was expected given these quantities’ strong correlations with each other.

## Discussion

In this work, we have shown that the commonly used measures of packing density, contact density (CN) and weighted contact density (WCN), capture non-local effects of amino-acid packing in a protein. To quantify the magnitude of these effects, we have defined purely local measures of packing density via the Voronoi tessellation, and we have shown that Voronoi cell volume encapsulates most of the relevant information about packing contained in various Voronoi cell characteristics. We have then shown that the variation in site-specific evolutionary rates (ER) can be partitioned into two independent components: variation explained by local, site-specific packing density (as measured by Voronoi cell volume) and variation explained by more long-range effects that extend beyond the first coordination shell around each site in a given protein. The longer-range effects play a non-negligible role in protein sequence evolution and explain on average approximately 10% of the observed variability in ER in the data set studied here. These 10% are in addition to and independent of the proportion of variance explained by the site-specific packing density, which explains on average another 34%. By contrast, the relative solvent accessibility (RSA) has negligible explanatory power for ER once site-specific packing is taken into account via Voronoi cell volume.

The quantities CN and WCN involve freely adjustable parameters in their definitions, and these parameters control to what extent CN and WCN assess the density of both immediate-neighbor and longer-range amino-acid packing. In principle, one could fine-tune these parameters for each individual protein to obtain the strongest correlations between ER and either CN or WCN^12^. Although the best parameterization of CN and WCN varies from one protein structure to another, we can take the values that maximize the median correlation for our dataset as the best overall parameterization of CN and WCN for our dataset. For both quantities, we find that the optimum parametrization is not entirely local. In particular, for WCN, which is the overall best predictor of ER in this dataset, the optimum exponent is −2.3, very close to the canonical value of −2 that is commonly used in the definition of WCN^11,12,20,21,39,41^. Since an exponent of −2 gives nearly equal weight to all shells of any radius around the focal residue, WCN is not really a site-specific measure of packing density. Instead it measures both the local packing density around the residue and the position of the residue relative to the global shape of the entire protein. By contrast, the Voronoi cell volume introduced here as an alternative measure of site-specific packing density depends exclusively on contributions from the nearest neighbors of each site in protein, that is, the first coordination shell around each focal residue.

Several earlier works^12,20^ have argued that local packing density is the best predictor of ER, while others^6,13^ have implicated RSA as the main determinant of sequence evolution, with local packing density playing a more peripheral role. Importantly, all these works have defined local packing density as CN or WCN. We have shown here that if we define local packing density as the inverse of the Voronoi cell volume, which by definition excludes the effects of longer-range effects beyond the first coordination shell, then it performs comparable to RSA. Our conclusion is that RSA and local packing density—when measured in a way that excludes longer-range effects—in principle represent the same characteristics of the local environment in a protein, that is, both quantities are proxy measures of the number of neighboring amino acids in the first coordination shell. Additional evolutionary variation is explained by non-local effects and/or the distribution of amino acids at larger distances, and these effects are captured by the most commonly used definition of WCN with inverse *r*^2^ weighting.

What is the origin of these non-local effects? There are two potential causes. First, longer-range effects can originate from steric interactions of local clusters of amino acids that extend beyond the first coordination shell. Evidence for such clusters comes from the discovery of protein sectors, regions of proteins with shared co-variation but little co-variation with other regions^47^. Further evidence comes from the observation that correlations between fluctuations of pairs of sites in proteins decay inversely with distances of the sites from each other^48^. Second, evolutionary variation may be influenced by shape and finite-size effects. If proteins have approximately uniform amino-acid density (as our work suggests, see next paragraph), then WCN measures primarily where in the three-dimensional protein structure a given residue is located. Thus, if there are systematic trends of evolutionary variation related to shape and finite size, for example conserved active sites that are located near the center of the protein, then WCN can partially capture these effects and provide further explanatory power for ER.

The finding that RSA and Voronoi cell volume are highly correlated was unexpected. According to their definitions, these two quantities measure different aspects of a residue’s local environment. In particular, RSA can only vary to the extent to which a residue is actually near the surface. For all completely buried residues RSA = 0, with no variability. By contrast, Voronoi cell volume depends on the exact location and number of amino-acid neighbors, and hence can, at least in principle, vary substantially inside the protein. The congruence of RSA and cell volume suggests that cell volume is approximately constant for buried residues and is determined primarily by solvent exposure on the protein surface. These conditions can only be satisfied if the amino-acid density is approximately constant throughout the protein. Thus, the strong correlations we have found between RSA and cell volume imply that amino-acid density is, to first order, uniform across protein structures.

One limitation of our present study is the choice of data set, 209 monomeric enzymes that have been previously studied by multiple groups. All enzymes in this data set are relatively small, globular, and have a well-defined active site, usually close to the center of the protein^28^. It is possible that the weighted contact number, with its radial weighting, performs particularly well on such proteins. For example, in the large and elongated influenza protein hemagglutinin, RSA explains over twice the variation in evolutionary rate than WCN does^15^. Moreover, active sites and protein–protein interfaces generally induce additional evolutionary constraints, and how these constraints interact with local packing density and longer-range steric interactions remains largely unexplored^6,15,49,50^. Therefore, future work will have to disentangle the contributions of local packing and longer-range effects in a wider set of protein structures, and it will also have to disentangle these contributions from the evolutionary constraints imposed by active sites and protein–protein interfaces.

## Appendix A Average Side-Chain Coordinates as the Best Representation of Protein 3D Structure

The calculations of the quantities CN and WCN, as well as the Voronoi tessellation, all require us to represent each amino acid by a reference point in 3D space. Most commonly, that reference point is chosen to be the C_*α*_ atom of the residue. However, this choice is mainly driven by convenience and convention, and *a priori* there is no particular reason to assume that this set of atomic coordinates is the best representation of individual sites in a protein. Indeed, earlier works have already suggested to use the center-of-mass of side chain coordinates to represent residues^51^. More recently, it was shown that WCN when calculated from side-chain center-of-mass coordinates generally results in significantly better correlations with ER than WCN calculated from *C*_*α*_ atoms does^21^.

To identify the best reference point in a more principled manner, we here considered seven different choices of atomic coordinates for the calculation of CN, WCN, and Voronoi cells. These include the set of coordinates of all backbone atoms (N, C, C_*α*_, O) and the first heavy atom in the amino acid side chains (C_*β*_). In addition, representative coordinates for each site in the protein can be also calculated by averaging over the coordinates of all heavy atoms in the side chains (referred to as SC). Finally, we calculated a representative coordinate by averaging over all heavy atom coordinates in both the side chain and the backbone of the amino acid (referred to as AA). For all calculations that required C_*β*_ coordinates, if the side chain *C*_*β*_ atom was not been resolved in the PDB file or the amino acid lacked *C*_*β*_ (i.e., Glycine), the *C*_*α*_ coordinate for the corresponding amino acid was used instead.

For all seven coordinate choices, we correlated both ER and RSA with both WCN and with Voronoi cell volume for each protein (Figure 5). In all cases, we found a systematic trend of declining correlation strengths as we moved the reference point further away from the geometric center of the side chain. In fact, the two atoms furthest away from the side chain, the nitrogen in the amino-group and the oxygen in the carboxyl group, generally produced the weakest correlations both with ER and with RSA. We also found that Voronoi cell volume and RSA measured approximately the same quantity, as long as the Voronoi tessellation was carried out on either SC or AA coordinates. The median correlation coefficient between RSA and cell volume for these coordinates was 0.92, and there was very little variation around that value (Figure 5D). (In comparison, the distribution of the correlation coefficients of side-chain WCN with cell volume had a median of 0.89, and on average, the correlations between cell volume and RSA were 0.05 higher than the correlations between cell volume and side-chain WCN.)

**Figure 5:**
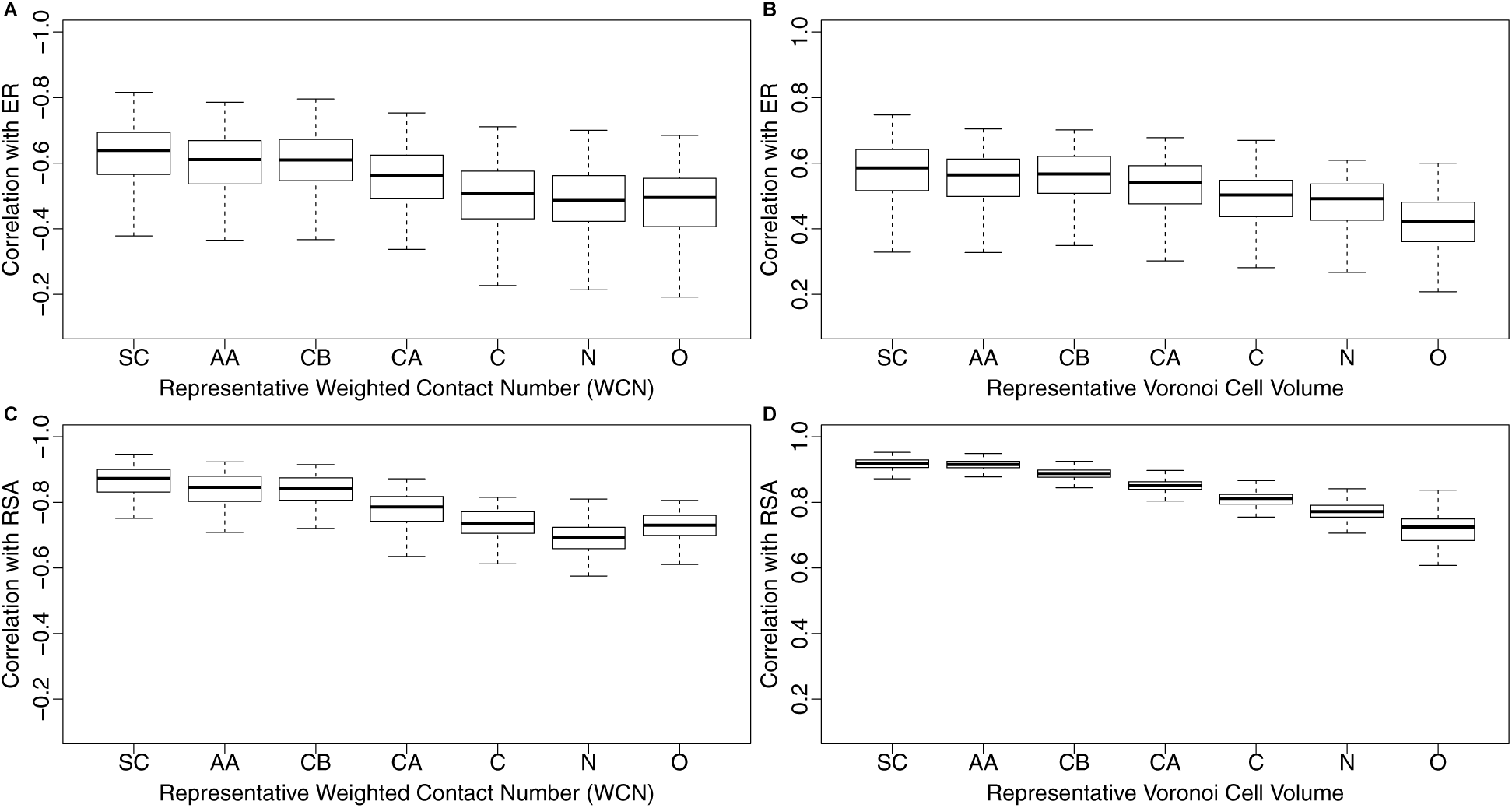
Side-chain centers provide most informative reference points for both WCN and Voronoi tessellation. (A) Distribution of correlations of WCN with ER, for seven different coordinate sets according to which WCN was calculated: SC, AA, CB, CA, N, C, O. Each coordinate set represents a different way of identifying the reference location of each residue. For SC (Side Chain) and AA (entire Amino Acid), the reference point is given, respectively, by the geometric average coordinates of the Side Chain (SC) atoms and the entire Amino Acid (AA) atoms. The latter include the backbone but exclude any Hydrogen. The coordinate sets CB, CA, N, C, and O use the respective atom in the amino acid as the reference point. (B) As in (A), but using Voronoi Cell Volume instead of WCN. (C) As in (A), but the correlations are calculated with RSA instead of with ER. (D) As in (B), but the correlations are calculated with RSA instead of with ER.

In combination, these results show that the biophysically relevant reference point for amino-acid position is the side chain, not the commonly used C_*α*_ atom. Further, they highlight that RSA properly accounts for the position of the side chain, and behaves nearly identically to side-chain Voronoi cell volume.

## Acknowledgements

We thank Julian Echave, Benjamin Jack, Austin Meyer, Stephanie Spielman, and Eleisha Jackson for helpful discussions and comments. This work was supported in part by NIH Grant R01 GM088344, DTRA Grant HDTRA1-12-C-0007, and the BEACON Center for the Study of Evolution in Action (NSF Cooperative Agreement DBI-0939454). The Texas Advanced Computing Center at UT Austin provided high-performance computing resources.

